# Polygenic risk for depression and resting state functional connectivity of subgenual anterior cingulate cortex in young adults

**DOI:** 10.1101/2024.02.18.580883

**Authors:** Yu Chen, Huey-Ting Li, Xingguang Luo, Guangfei Li, Jaime S. Ide, Chiang-Shan R. Li

## Abstract

Genetic variants may confer risks for depression by modulating brain structure and function. Prior evidence has underscored a key role of the subgenual anterior cingulate cortex (sgACC) in depression. Here, we built on the literature and examined how the resting state functional connectivity (rsFC) of the sgACC was associated with polygenic risks for depression. We followed published routines and computed seed-based whole-brain sgACC rsFC and polygenic risk scores (PRS) of 717 young adults curated from the Human Connectome Project. We performed whole-brain regression against PRS and severity of depression symptoms in a single model for all subjects and for men and women alone, controlling for age, sex (for all), race, severity of alcohol use, and household income, and evaluated the results at a corrected threshold. We found lower sgACC rsFC with the default mode network and frontal regions in association with PRS and lower sgACC-cerebellar rsFC in association with depression severity. We also noted sex differences in the connectivity correlates of PRS and depression severity. In an additional set of analyses, we observed a significant correlation between PRS and somatic complaints score and altered sgACC-somatosensory cortical connectivity in link with the severity of somatic complaints. Our findings collectively highlighted the pivotal role of distinct sgACC-based networks in the genetic predisposition to depression and the clinical manifestation of depression. Distinguishing the risk from severity markers of depression may have implications in developing early and effective treatments for individuals at risk for depression.

## 1. Introduction

Depression is a heritable psychiatric disorder and a leading cause of disability and suicide. Identifying the biomarkers has long been a focus in clinical neuroscience and therapeutics research of depression. Converging evidence has associated depression with brain network dysfunction and highlighted the role of frontolimbic circuits in the pathophysiology of depression. For instance, relative to healthy controls, both adolescents and adults with depression showed altered brain morphometry and connectivity of the ventromedial frontal cortex (vmPFC), anterior cingulate cortex (ACC), and subcortical structures, including the striatum and amygdala (Kaiser et al., 2015; Kerestes et al., 2014; Koolschijn et al., 2009; Mulders et al., 2015; Sacher et al., 2012; Schmaal et al., 2017).

In particular, as a hub of the emotion regulation circuit (Scharnowski et al., 2020), the subgenual ACC (sgACC) is widely implicated in depression, showing both structural and functional changes in patients with major depressive disorder (MDD; Gotlib and Hamilton, 2008). Specifically, people with MDD vs. controls showed lower gray matter volumes (GMV) of the sgACC (Rodriguez-Cano et al., 2014) and stronger resting state functional connectivity (rsFC) of the sgACC with insula, amygdala, and hippocampus as well as weaker rsFC with the precuneus (Connolly et al., 2013; de Kwaasteniet et al., 2013), a core structure of the default mode network (DMN). In people with MDD, these volumetric and connectivity changes were associated with depression severity (Connolly et al., 2013; de Kwaasteniet et al., 2013; Jaworska et al., 2016; Rodriguez-Cano et al., 2014). Further, treatment studies showed that brain stimulation targeting the sgACC appeared to demonstrate superior efficacy as compared to other regions of interest (Drobisz and Damborska, 2019; Schlaepfer and Bewernick, 2013). Together, these studies provide a substantial body of evidence supporting a critical role of the sgACC in the pathophysiology of depression.

The great majority of these studies were conducted in patients with depression and oftentimes chronic depression, and it remains unclear whether or to what extent the findings reflect the risks vs. consequences of depression. Earlier studies suggested a genetic basis of sgACC dysfunction in depression. For instance, carriers of the short allele of a functional 5’ promoter polymorphism of the serotonin transporter gene (5HTTLPR) showed elevated risk of depression (Lotrich and Pollock, 2004). Importantly, a multimodal imaging study identified an effect of 5-HTTLPR genotype on sgACC structure and functional connectivity. As compared to long/long allele genotype, short-allele carriers showed significantly reduced sgACC GMV and enhanced sgACC-amygdala coupling, in link with the risk for depression (Pezawas et al., 2005).

Although genome-wide association studies (GWAS) have identified many genetic, including 5HTTLPR, variants of depression (Flint and Kendler, 2014; Howard et al., 2019; McIntosh et al., 2019; Wray et al., 2018), each variant confers only a small effect in the risk for depression. Investigators have thus employed the polygenic risk score (PRS) to capture the overall individual genetic risks for depression (Cao et al., 2021; Halldorsdottir et al., 2019). However, to date, it remains unclear whether the sgACC markers may underpin the overall genetic risks for depression.

The current study aimed to examine how the sgACC rsFC manifested in the overall genetic risks for depression and whether these markers could be distinguished from those reflecting the severity of depression symptoms. To this end, we employed the clinical, resting state fMRI, and genotyping data of a large sample of young adults of the Human Connectome Project (HCP). We computed the PRS leveraging the meta-analysis of GWAS of 33 UK Biobank cohorts of the Psychiatric Genomics Consortium as the base data. We performed whole-brain voxel-wise regressions on the seed-based sgACC rsFC against PRS and symptom severity score in the same model to identify the connectivity correlates each of genetic risk and severity of depression problems. We also investigated the connectivity correlates of somatic complaints, which, amongst the symptoms of depression, showed the strongest correlation with the PRS for depression. We also performed the regressions separately in men and women and investigated sex differences with slope tests. We broadly hypothesized that sgACC rsFC with the frontolimbic network would reflect the genetic risks for depression and that the genetic risks and severity of depression problems would involve distinct sgACC connectivities.

## 2. Methods

### 2.1 Dataset: subjects and assessments

The HCP Young Adult (HCP-YA) S1200 release contains clinical, behavioral, and 3T magnetic resonance (MR) imaging data of 1,206 young healthy adults without severe neurodevelopmental, neuropsychiatric, or neurologic disorders. A total of 489 subjects were excluded due to questionable image quality or poor image segmentation (n = 213), missing rsfMRI, cardiac/respiratory, or clinical data (n = 85), or excessive head movements during rsfMRI (n = 191; see details in “*2.2 MRI data acquisition and preprocessing*”). Thus, the data of a final sample of 717 participants (377 women, age 22-36 years) were included in the analyses.

Participants completed the Achenbach Adult Self Report (ASR; Achenbach et al., 2003), where individual questions were rated on a 3-point Likert scale (0-Not True, 1-Somewhat or Sometimes True, 2-Very True or Often True), including the DSM-oriented subscale of depression (14 items). In addition, the ASR included 13 other subscales that may be related to the clinical manifestation of depression, e.g., somatic complaints (12 items). The age-and sex-adjusted *T* scores of depression and other subscales were used in the analyses, with higher *T* scores indicating higher severity of depression and other symptoms. The DSM-oriented depression severity *T* score was used as the target phenotype for the computation of PRS for depression. Participants were also evaluated with the Semi-Structured Assessment for the Genetics of Alcoholism (SSAGA; Bucholz et al., 1994). As in our previous studies (Li et al., 2022), we performed a principal component analysis on the 15 interrelated drinking metrics and identified one principal component (PC1) with an eigenvalue > 1 that accounted for 50.54% of the variance. Higher PC1 values indicate more severe drinking. We controlled for the effects of alcohol use by including drinking PC1 as a covariate in all analyses, since depression and alcohol misuse frequently comorbid (Chen et al., 2022b). SSAGA also collected household income data for each subject: <$10,000 = 1,10K-19,999 = 2, 20K-29,999 = 3,30K-39,999 = 4, 40K-49,999 = 5, 50K-74,999 = 6, 75K-99,999 = 7, ≥100,000 = 8. Prior evidence has shown an inverse relation between household income and depression (Akhtar-Danesh and Landeen, 2007; Hounkpatin et al., 2015). Therefore, we included the household income as an additional covariate in all analyses.

We performed independent sample *t* tests on age, drinking PC1, and household income and a chi-square test on race to examine sex differences. These variables were included in all the following analyses as covariates of men and women combined and (except for sex) separately.

We performed independent sample *t* tests on the PRS, and depression and somatic complaints *T* scores to examine sex differences. We conducted partial correlations of the PRS with depression and somatic complaint *T* scores in men and women combined and separately, with ancestry proportions as an additional set of covariates. We used slope tests to examine sex differences in these correlations (Zar, 1999).

### 2.2 MRI data acquisition and preprocessing

As described in our previous studies (Chen et al., 2022a; Chen et al., 2021), the details of the data collection procedures can be found in the HCP S1200 Release Reference Manual. All imaging data were acquired on a customized Siemens 3T Skyra with a standard 32-channel Siemens receiver head coil and a body transmission coil. T1-weighted high-resolution structural images were acquired using a 3D MPRAGE sequence with 0.7 mm isotropic resolution (FOV = 224 mm, matrix = 320, 256 sagittal slices, TR = 2400 ms, TE = 2.14 ms, TI = 1000 ms, FA = 8°) and used to register rsfMRI data to a standard brain space. The rsfMRI data were collected in two sessions, using gradient-echo echo-planar imaging (EPI) with 2.0 mm isotropic resolution (FOV = 208 ×180 mm, matrix = 104 ×90, 72 slices, TR = 720 ms, TE = 33.1 ms, FA = 52°, multi-band factor = 8). Within each session, oblique axial acquisitions alternated between phase encoding in a right-to-left (RL) direction in one run and phase encoding in a left-to-right (LR) direction in the other run. Each run lasted 14.4 minutes (1200 frames). Physiological data (i.e., cardiac and respiratory signals) associated with each functional MR scan were also acquired, using a standard Siemens pulse oximeter placed on a digit and a respiratory belt placed on the abdomen. These physiological signals were sampled equally at 400 Hz (∼288 samples per frame).

We processed the first session of the rsfMRI data with Statistical Parametric Mapping (SPM8). Images of each participant were first realigned (motion corrected) and a mean functional image volume was constructed from the realigned image volumes. These mean images were co-registered with the high-resolution structural MPRAGE image and then segmented for normalization with affine registration followed by nonlinear transformation. The normalization parameters determined for the structural volume were then applied to the corresponding functional image volumes for each participant. Afterwards, the images were smoothed with a Gaussian kernel of 4 mm at Full Width at Half Maximum.

White matter and cerebrospinal fluid signals, whole-brain mean signal, and physiological signals were regressed out to reduce spurious BOLD variances and to eliminate cardiac-and respiratory-related artifacts. A temporal band-pass filter (0.009 Hz < *f* < 0.08 Hz) was also applied to the time course to obtain low-frequency fluctuations, as in our previous studies (Li et al., 2023; Zhang et al., 2021; Zhu et al., 2022; Zhu et al., 2023). Lastly, to eliminate global motion-related artifacts, a “scrubbing” method was applied (Dong et al., 2023). Specifically, frame-wise displacement given by *FD*(*t*) = |Δ*d*_x_(*t*)| + |Δ*d*_y_(*t*)| + |Δ*d*_z_(*t*)| + |Δ*α*(*t*)| + |Δ*β*(*t*)| + |Δ*γ*(*t*)| was computed for every time point *t*, where (*d*_x_, *d*_y_, *d*_z_) and (*α*, *β*, *γ*) are the translational and rotational movements, respectively (Power et al., 2012). Moreover, the root mean square variance of the differences (DVARS) in % BOLD intensity *I*(*t*) between consecutive time points across brain voxels, was computed as: *DVARS*(*t*) = sqrt(|*I*(*t*) – *I*(*t*-1)|^2^), where the brackets indicate the mean across brain voxels. Following previous HCP studies (Chen and Li, 2023; Li et al., 2019), we marked volumes with FD > 0.2mm or DVARS > 75 as well as one frame before and two frames after these volumes as outliers (censored frames). Uncensored segments of data lasting fewer than five contiguous volumes were also labeled as censored frames. A total of 191 participants who had either run with more than half of the frames flagged as censored were removed from further analyses.

### 2.3 sgACC rsFC computation

Whole-brain voxel-wise analyses were conducted to compute the seed-based rsFC of the sgACC. We used a mask of the sgACC from the Automated Anatomic Labeling atlas (Tzourio-Mazoyer et al., 2002). The BOLD time courses of each voxel of the sgACC seed were averaged, and the correlation coefficients were computed between the average time course and the time courses of all other voxels for individual participants. The correlation matrix was Fisher’s *z*-transformed into *z* score maps and averaged across the two runs for each participant.

### 2.4 Genotyping and PRS computation

The base sample included 170,756 unrelated cases with MDD and 329,443 unrelated healthy controls from the meta-analysis of GWAS of 33 UK Biobank cohorts of the Psychiatric Genomics Consortium (PGC; Howard et al., 2018; Hyde et al., 2016; Wray et al., 2018). MDD cases were diagnosed according to DSM-IV. The target HCP-YA sample included 453 families with 1,140 healthy twins/siblings (618 women). The depression *T* score served as the phenotype in PRS analysis. Accurate computation of the PRS requires a large sample; thus, we included all subjects with available genotype data in the computation of PRS.

#### 2.4.1 Quality control for both base and target samples

The base sample was genotyped on microarrays and then 8,483,301 single nucleotide polymorphisms (SNPs) were imputed. The HCP-YA sample was genotyped using Illumina Infinium Multi-Ethnic Genotyping Array (MEGA) or Infinium Neuro Consortium Array (2,292,654 SNPs). The computation of PRS requires base (summary statistics of GWAS), target (whole-genome genotype and phenotype), and covariate (age, sex, and principal components of ancestry) data (Choi et al., 2020). Both base and target data underwent quality control based on a standard protocol (Choi et al., 2020; Zuo et al., 2012), to filter out the SNPs with ambiguous alleles or those mismatching between base and target data, and to exclude those with sexes mismatching between self-report and sex chromosomes. We also removed the subjects with genetic relatedness for any pair of individuals between base and target samples. The final base data had a chip-heritability > 0.05, and the risk alleles all had log(OR) scores > 0 that reflected the effect size of SNP-MDD associations. Only the SNPs existing in both “cleaned” base and target samples were included for the computation of PRS.

#### 2.4.2 PRS computation

PRS was computed according to a formula (Choi et al., 2020) that considered the effect size of SNPs, the number of risk alleles, and the total number of SNPs included (based on different *p* values). In brief, PRS was computed as a sum of the genome-wide risk alleles, weighted by the corresponding effect size estimated from GWAS. We calculated the PRS for each subject using the program PRSice-2 (Choi et al., 2020), which automatically excluded the correlated SNPs (pairwise *r*^2^ > 0.25) and considered age, sex (when mixing males and females), and principal components of ancestry as covariates. We derived the “best-fit” PRS from different *p* value thresholds and the percentage of phenotypic variation explained by the “best-fit” PRS. The “best-fit” PRS with the best discriminative capacity was determined based on the maximal area under the receiver-operator curve in a regression model with the phenotype as the outcome and the PRS, age, sex (when computed for males and females together), and ancestry proportions as covariates (Khera et al., 2018).

To keep the individuals unrelated within each target dataset, as required to ensure the accuracy of PRS, the twins and siblings within each family in HCP-YA samples were randomly separated into independent target subsets. To maximize the sample size of each subset, the subjects within the first subset who were unrelated to the subjects within other subsets were filled in each of the latter subsets repeatedly. Then, the PRS and the percentage of phenotypic variation explained by the PRS (i.e., R^2^) were calculated separately for each subset under the same *p*-value thresholds. The PRS scores of the same subject, if showing < 1% difference across different subsets, were averaged as the final PRS. And the percentages of phenotypic variation, if showing < 1% difference across different subsets, were averaged across the subsets as the final values.

### 2.5 Behavioral, whole-brain regression and region-of-interest (ROI) analyses

We first performed whole-brain linear regressions against PRS and depression *T* scores in the same model. We evaluated the results with uncorrected voxel *p* < 0.001 in combination with cluster *p* < 0.05 corrected for family-wise error (FWE) of multiple comparisons on the basis of Gaussian random field theory as implemented in the SPM, with 8 voxels expected per cluster.

In ROI analyses, for all significant clusters identified in whole-brain regressions in men and women separately, we estimated *β* values of sgACC rsFC and computed the partial correlations of sgACC rsFC with depression *T* score and PRS in men and women separately and followed up with slope tests to examine sex differences in the correlations. Note that the slope tests of sex differences did not represent “double-dipping” (Chen et al., 2022c), as the regression maps were identified with a threshold and clusters showing significant correlations in men might have just missed the threshold in women, and vice versa. Thus, direct tests of the slopes were needed to confirm sex differences.

We observed that the PRS for depression was significantly correlated with somatic complaints *T* score (see Results 3.1). Thus, in an additional set of analyses, we included PRS, depression *T* score, and somatic complaints *T* score in the same whole-brain regression model, with the same set of covariates, in men and women together and separately. We also estimated *β* values of sgACC rsFC for any clusters identified in whole-brain regressions in men and women alone and computed the partial correlations of sgACC rsFC with somatic complaints *T* score for men and for women, with slope tests to investigate sex differences in the correlations.

## 3. Results

### 3.1 Demographic and clinical measures

The descriptive statistics and *t* or chi-square tests of sex differences are shown in **Table 1**. Men were significantly older and had greater drinking severity (*p*’s < 0.001). Men and women did not show significant differences in race composition (*p* = 0.779) or household income (*p* = 0.089). With age, race, drinking PC1, and household income as covariates, men vs. women did not show any significant differences in the PRS (*p* = 0.969), depression *T* score (*p* = 0.650), or somatic complaints *T* score (*p* = 0.962).

**Table 1.** Demographic and clinical measures of men and women.

| | Men (n = 340) | Women (n = 377) | $t/\chi^2$ | $p$ |
| --- | --- | --- | --- | --- |
| Age (years) | 27.54 ± 3.55 | 29.35 ± 3.67 | -6.70 | **< 0.001** |
| Race |  |  | 2.48 | 0.779 |
| AI/AN | 1 (0.29%) | 0 (0%) |  |  |
| A/NH/OPI | 23 (6.76%) | 23 (6.10%) |  |  |
| AA | 35 (10.30%) | 48 (12.73%) |  |  |
| White | 267 (78.53%) | 290 (76.92%) |  |  |
| > one race | 10 (2.94%) | 10 (2.65%) |  |  |
| UK/NR | 4 (1.18%) | 6 (1.60%) |  |  |
| Drinking PC1 | 0.36 ± 1.08 | -0.33 ± 0.79 | 9.73 | **< 0.001** |
| Household income | 5.01 ± 2.18 | 5.28 ± 2.14 | -1.70 | 0.089 |
| PRS | 0.0109080 ± 0.0000423 | 0.0191068 ± 0.0000417 | 0.04 | 0.969 |
| Depression $T$ score | 53.93 ± 5.90 | 53.70 ± 5.33 | -0.45 | 0.650 |
| Somatic complaints $T$ score | 53.88 ± 5.69 | 53.58 ± 5.51 | 0.05 | 0.962 |

*Note:* Values are mean ± SD or n (%). AI/AN: American Indian/Alaskan Native; A/NH/OPI: Asian/Native Hawaiian/Other Pacific Islander; AA: African American; UK/NR: Unknown/Not Reported; PC1: first component of drinking metrics; PRS: polygenic risk score. The independent $t$ tests for PRS, depression $T$ score, and somatic complaints $T$ score were performed with age, race, drinking PC1, and household income as covariates. Bold $p$ values indicate significant sex differences.

For the entire cohort with genotype data (n = 1,140; 618 women) in the HCP, greater PRS was associated with higher depression *T* score, with age, sex (for all subjects), race, drinking PC1, household and ancestry proportions as covariates (*r* = 0.07, *p* = 0.019; **Figure 1A**). Men also showed a significant correlation between PRS and depression *T* score (*r* = 0.12, *p* = 0.009) but not women (*r* = 0.02, *p* = 0.624). However, the slope of regression did not differ significantly between men and women (*t* = 1.68, *p* = 0.093). In the current, smaller imaging sample (n = 717; 377 women), greater PRS was significantly correlated with higher depression *T* score in men (*r* = 0.12, *p* = 0.041) but not in women (*r* = −0.020, *p* = 0.713) or across all subjects (*r* = 0.06, *p* = 0.133), with age, sex (for all subjects), race, drinking PC1, household and ancestry proportions as covariates. The slope test did not show significant sex difference in the correlations (*t* = 1.63, *p =* 0.103).

**Figure 1.**
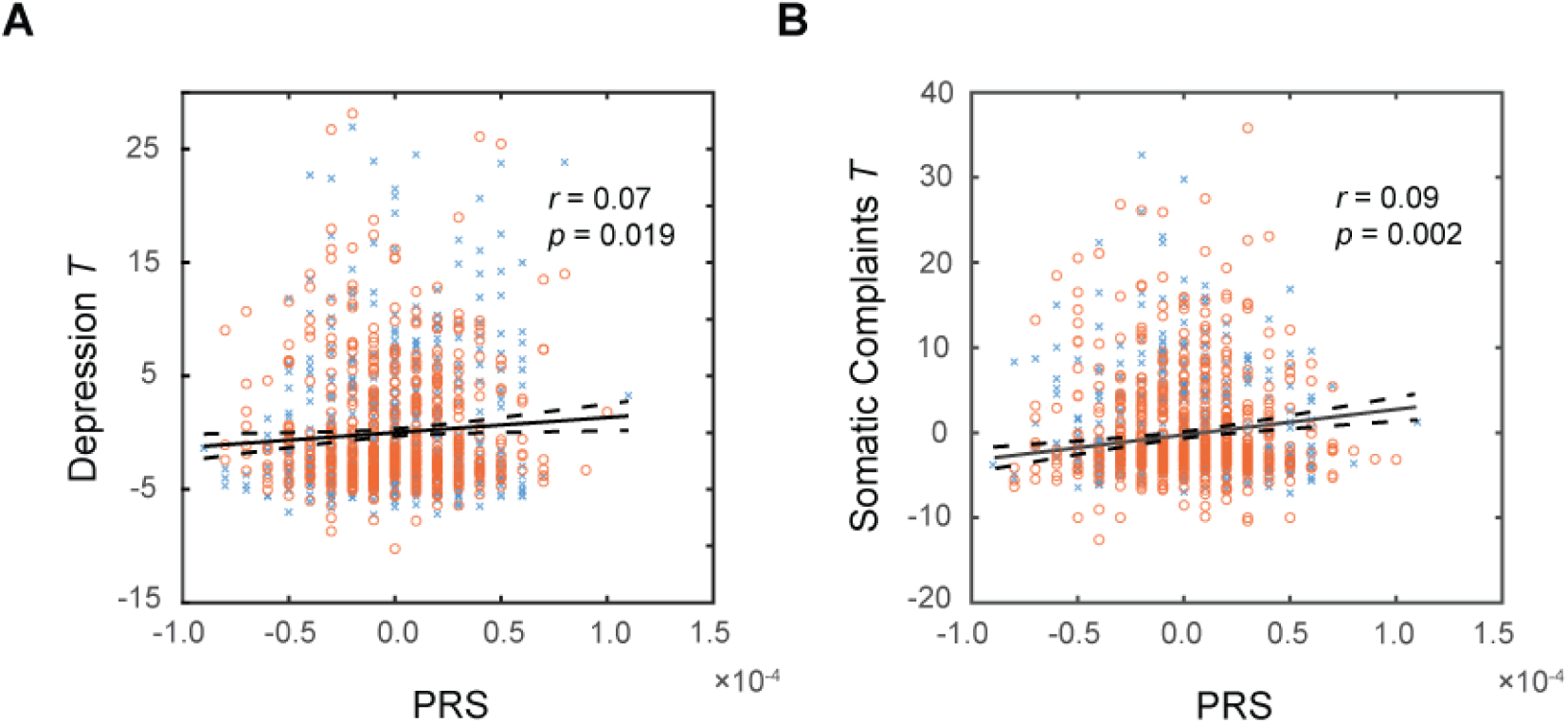
Scatter plots showing the partial correlations of PRS with **(A)** depression *T* score, and **(B)** somatic complaints *T* score, in 1,140 subjects (618 women) with age, sex (for all subjects), race, drinking PC1, household income and ancestry proportions as covariates. Note that residuals are presented here (blue crosses: men; orange circles: women). Solid and dashed lines represent the regressions and 95% confidence intervals, respectively. The coefficient *r* and *p* values are also presented.

For the entire cohort with genotype data, greater PRS was also associated with higher somatic complaint *T* score, with age, sex (for all subjects), race, drinking PC1, household and ancestry proportions as covariates (*r* = 0.09, *p* = 0.002; **Figure 1B**), with correction for multiple comparisons (*p* < 0.05/14 = 0.0036). Men (*r* = 0.09, *p* = 0.041) and women (*r* = 0.09, *p* = 0.023) alone also showed a significant correlation between PRS and somatic complaints *T* score, with the slope test *t* = 0.65 and *p* = 0.513. In the current smaller imaging sample, however, PRS was not significantly correlated with somatic complaints *T* score in men (*r* = 0.05, *p* = 0.359) or women (*r* = 0.04, *p* = 0.442) alone or across all subjects (*r* = 0.06, *p* = 0.132), with the same covariates.

### 3.2 Seed-based sgACC rsFC in relation to PRS and depression T score

The whole-brain sgACC rsFCs in correlation with PRS and depression *T* score are shown in **Figure 2** and the clusters are summarized in **Supplementary Table S1**. In all subjects (**Figure 2A**), higher PRS were correlated with weaker sgACC rsFC with bilateral superior frontal gyri (SFG) and bilateral posterior cingulate cortex/ventral precuneus (PCC/vPCu), and higher depression *T* scores were correlated with weaker rsFC between sgACC and right cerebellum (CBL). In men alone (**Figure 2B**), higher PRS were correlated with stronger sgACC-left CBL and weaker sgACC-left SFG rsFC, and higher depression *T* scores were correlated with weaker sgACC rsFC with bilateral CBL and insula (INS). In women alone (**Figure 2C**), higher PRS were correlated with stronger sgACC rsFC with bilateral lingual gyri and calcarine sulcus (LING/CAL).

**Figure 2.**
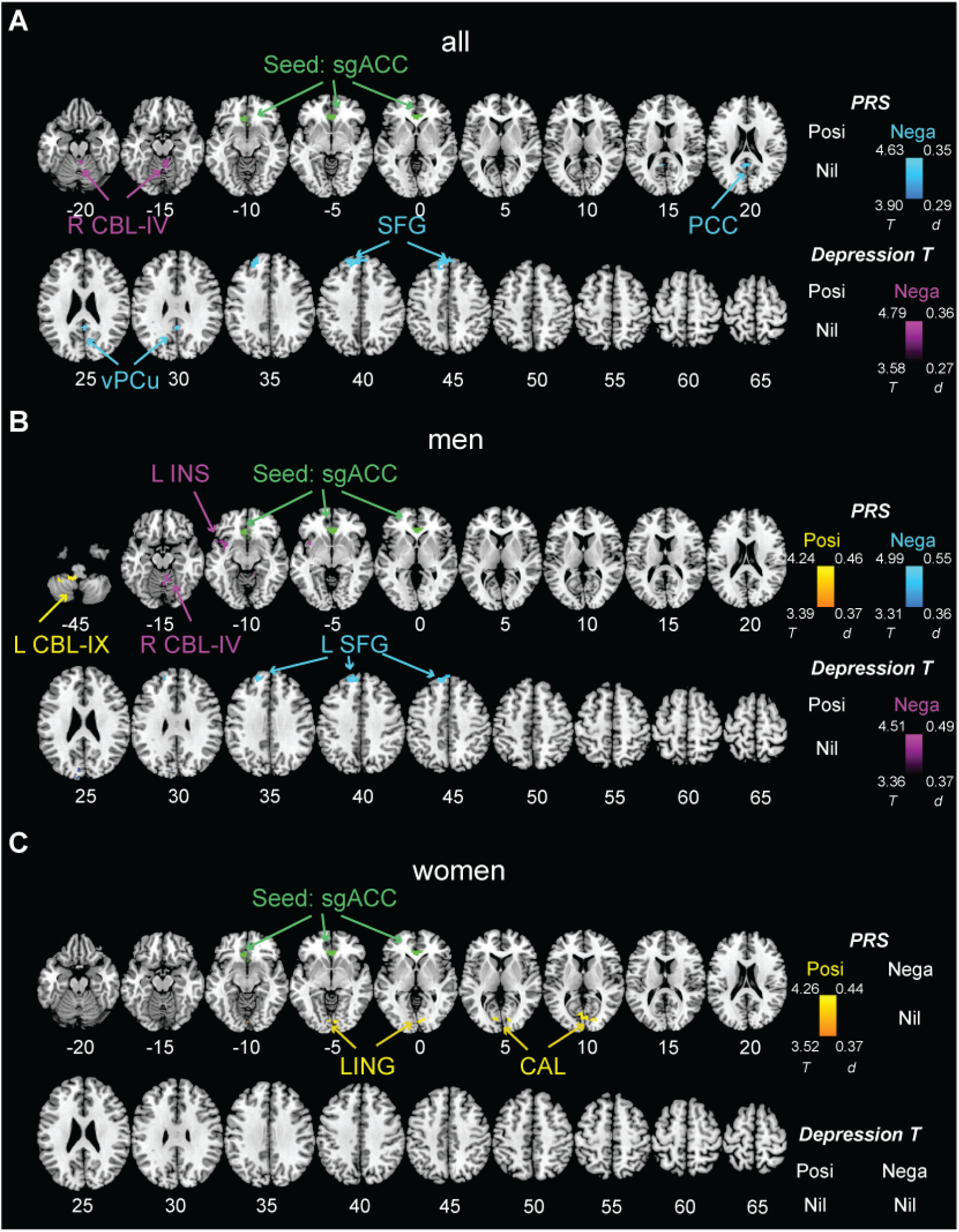
Seed-based whole-brain rsFC of sgACC (green) in correlation with polygenic risk score (PRS) and depression *T* score for **(A)** all subjects, **(B)** men, and **(C)** women. PRS and depression *T* score were modeled together, with age, sex (for all subjects), race, drinking principal component (PC1), and household income as covariates. The results were evaluated at voxel *p* < 0.001, uncorrected in combination with cluster *p* < 0.05 family-wise error (FWE) corrected. Warm color: in positive (Posi) correlation with PRS; Cool color: in negative correlation with PRS; Violet: in negative (Nega) correlation with depression *T* score; Color bar shows voxel *T* and Cohen’s *d* values. L: left; R: right; CBL: cerebellum; sgACC: subgenual anterior cingulate cortex; PCC: posterior cingulate cortex; vPCu: ventral precuneus; SFG: superior frontal gyrus; INS: insula; LING: lingual gyrus; CAL: calcarine sulcus.

For the 5 clusters identified in men and women alone with sgACC rsFCs showing significant correlations with PRS or depression *T* score, we confirmed sex differences in the correlations with slope tests. Considering correction for multiple comparisons (*p* < 0.05/5 = 0.01), men vs. women showed significantly stronger correlations between PRS and sgACC rsFC with left cerebellum lobule IX and between depression *T* score and sgACC rsFC with left insula and right cerebellum lobule IV, as well as weaker correlations between PRS and sgACC rsFC with lingual gyrus and calcarine sulcus (**Figure 3**), with the slope test *t’s* ≥ 2.40 and *p’s* ≤ 0.001.

**Figure 3.**
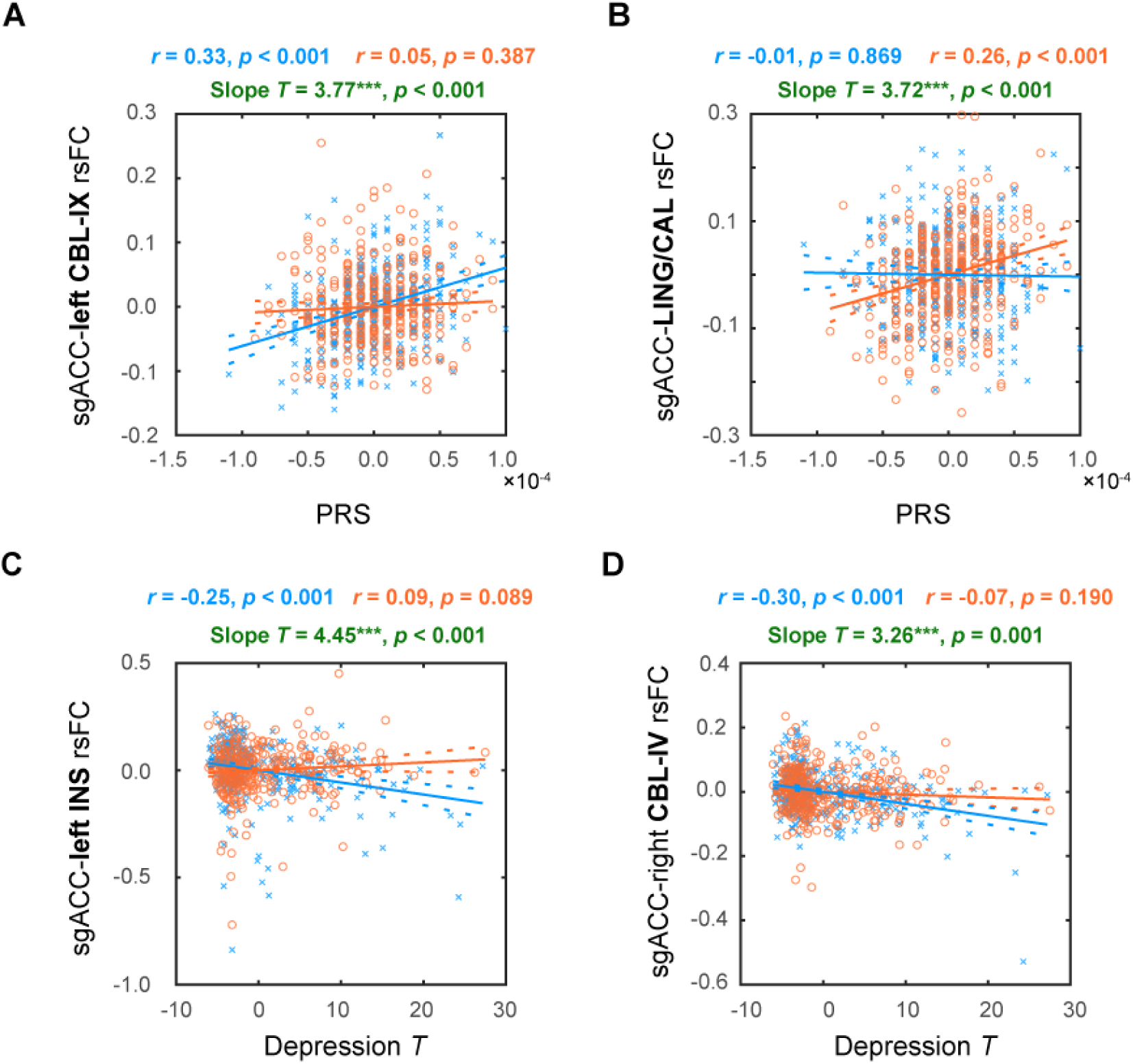
Scatter plots showing sex differences in the correlations of sgACC rsFC with PRS and depression *T* score. **(A)** PRS vs. sgACC-left CBL-IX rsFC; **(B)** PRS vs. sgACC-LING/CAL rsFC; **(C)** Depression *T* vs. sgACC-left INS rsFC; and **(D)** Depression *T* vs. sgACC-right CBL-IV rsFC. The data points represent residuals (men: in blue; women: in orange). Solid and dashed lines represent the regressions and 95% confidence intervals, respectively. The coefficients *r* and *p* values for the correlations as well as slope *t* and *p* values for sex differences in the correlations are presented on top of each panel. Slope test \*\*\**p* < 0.05/5 = 0.01 for multiple comparisons. *Note*: sgACC: subgenual anterior cingulate cortex; rsFC: resting state functional connectivity; PRS: polygenic risk score; CBL: cerebellum; LING: lingual gyrus; CAL: calcarine sulcus; INS: insula.

### 3.3 Seed-based sgACC rsFC in relation to somatic complaints T score

We included the PRS, depression *T* score, and somatic complaints *T* score as regressors with the same covariates in an additional model. The whole-brain sgACC rsFCs in correlation with somatic complaints *T* score are shown in **Figure 4** and the clusters are summarized in **Supplementary Table S2**. In all subjects (**Figure 4A**), higher scores of somatic complaints were correlated with weaker sgACC rsFC with a cluster in the left postcentral and supramarginal gyri. In men alone (**Figure 4B**), we did not observe any significant clusters. In women alone (**Figure 4C**), higher scores of somatic complaints were correlated with lower sgACC-dorsal precuneus (dPCu) rsFC. The slope test confirmed that women (*r* = −0.14, *p* = 0.009) vs. men (*r* = 0.04, *p* = 0.498) showed stronger correlations between sgACC-dPCu rsFC and somatic complaints *T* score (*t* = 2.34, *p* = 0.019).

**Figure 4.**
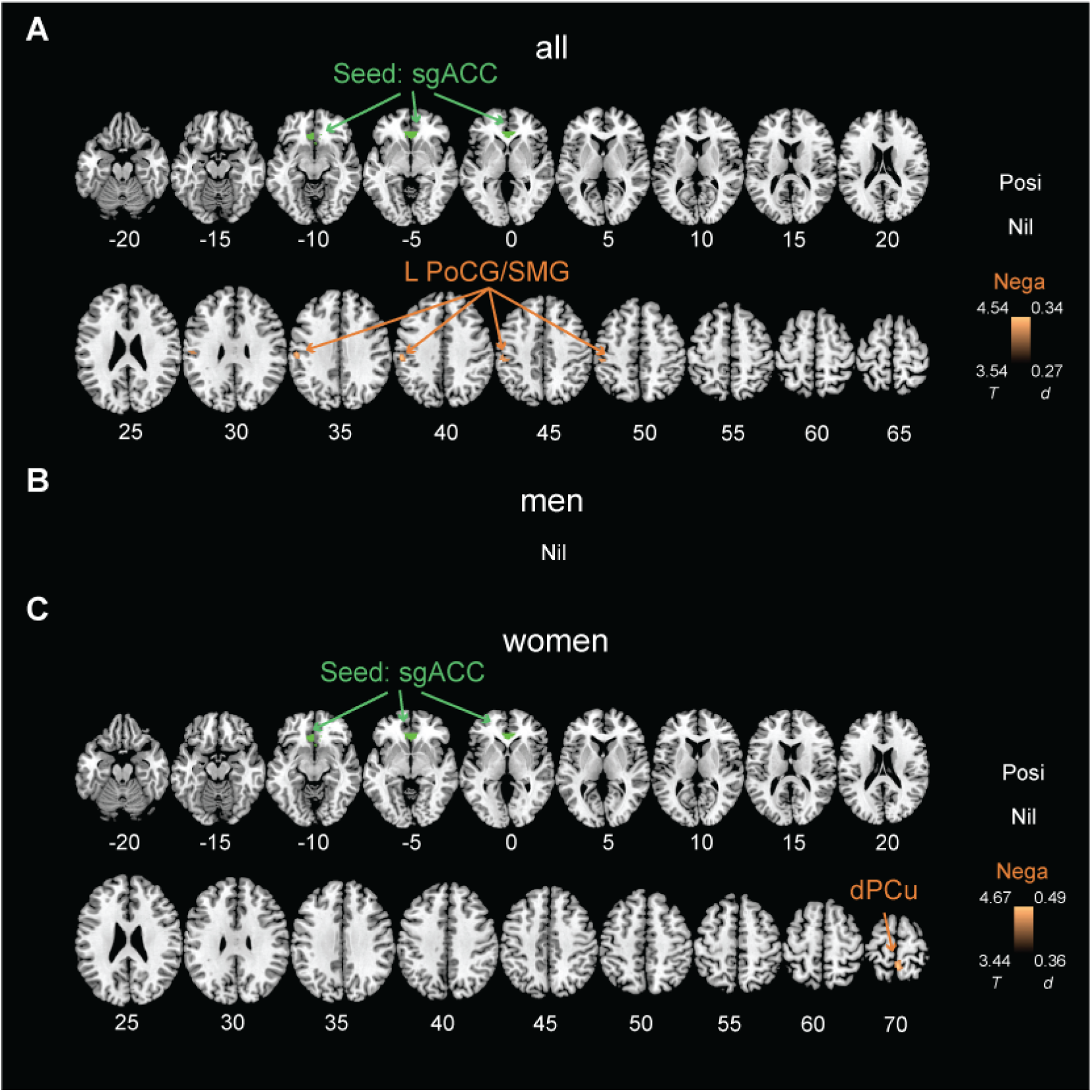
Seed-based whole-brain rsFC of sgACC (green) in correlation with somatic complaints *T* score for **(A)** all subjects, **(B)** men, and **(C)** women. The somatic complaints *T* score was modeled together with PRS and depression *T* score, controlling for age, sex (for all subjects), race, drinking principal component (PC1), and household income as covariates. The results were evaluated at voxel *p* < 0.001, uncorrected in combination with cluster *p* < 0.05 family-wise error (FWE) corrected. The results for PRS and depression *T* score is shown in the **Supplementary Figure S1**. Nil: no significant findings; Brown: in negative (Nega) correlation with somatic complaints *T* score; Color bar shows voxel *T* and Cohen’s *d* values. L: left; sgACC: subgenual anterior cingulate cortex; PoCG: postcentral gyrus; SMG: supramarginal gyrus; dPCu: dorsal precuneus.

In this model, the sgACC rsFC correlates of PRS and depression *T* score remained largely unchanged (see **Supplementary Figure S1** and **Table S2**), except that a few new clusters could be identified. Specifically, in men, sgACC rsFC with left hippocampus was negatively correlated with the depression *T* score. In women, sgACC rsFC with right middle temporal gyrus was positively correlated with PRS, and sgACC rsFC with right calcarine sulcus and with right supplementary motor area (SMA) was negatively and positively correlated with the depression *T* score, respectively.

## 4. Discussion

We combined genetic analysis, neuroimaging, and psychological assessments to delineate seed-based sgACC connectivity markers that underlie genetic risks for depression and clinical manifestation of depression. We observed distinct functional connectivities of the sgACC each in relation to PRS and to the severity of depression symptoms in a large sample of young adults. Across all subjects, higher genetic susceptibility to depression was associated with lower functional coupling of the sgACC with the DMN and superior frontal cortex, whereas greater severity of depression symptoms was associated with lower sgACC-right cerebellum connectivity. Moreover, the severity of somatic complaints was associated with reduced sgACC connectivity with a cluster encompassing the left postcentral and supramarginal gyri (PoCG/SMG). We also identified sgACC connectivities in relation to genetic risks, depression, and somatic complaints in men and women separately. Specifically, men (vs. women) and women (vs. men) each showed significantly higher sgACC rsFC with the left CBL-IX (posterior cerebellum) and with bilateral LING/CAL, respectively, in correlation with PRS. Men vs. women also showed significantly lower sgACC rsFC with the left INS and bilateral CBL-IV (anterior cerebellum) in correlation with depression *T* score. Women vs. men presented stronger negative correlation between sgACC-dPCu rsFC and somatic complaints. Together, these findings distinguished risk-and severity-related neural markers of depression and highlighted sex differences in these connectivity markers. We discussed the main findings below.

### 4.1 sgACC rsFC underlying genetic risks and severity of depression

We observed negative correlations between PRS for depression and the functional coupling of the sgACC with a cluster encompassing the ventral precuneus (vPCu) and PCC and another cluster in the left/medial SFG, controlling for the severity of depression. These findings suggest that prior report of diminished sgACC with the precuneus and frontal cortical regions in patients with MDD may largely reflect the cumulative genetic risks of depression (Connolly et al., 2013). The precuneus, particularly the vPCu (Zhang and Li, 2012), and PCC are both core regions of the DMN, a network that supports self-referential activities (Gusnard et al., 2001) and a myriad of cognitive (Spreng et al., 2009) and emotional (Satpute and Lindquist, 2019) functions. DMN dysfunction is implicated in rumination and preoccupation with negative information in patients with MDD as well as individuals at risk for depression (Chou et al., 2023). Other studies implicated the sgACC and SFG in emotion regulation (Drevets et al., 2008; Frank et al., 2014), and this circuit dysfunction conduces to the development of affective disorders (Savitz and Drevets, 2009; Whalley et al., 2012).

We also found reduced sgACC rsFCs with the right cerebellum lobule IV in correlation with more severe depression symptoms. Though best known for its role in motor control, the cerebellum is central to cognitive and emotion processing (Buckner, 2013; Depping et al., 2018; Fastenrath et al., 2022; Schmahmann, 2019; Schutter and Van Honk, 2005). Studies have accumulated to support altered cerebello-striato-prefrontal network function in people with depressive disorders (Depping et al., 2018; Lupo et al., 2019). It is suggested that disruptions of prefrontal and limbic structures and functions are present in the early stages of mood disorders, whereas cerebellar dysfunction may reflect a consequence of these disorders (Lupo et al., 2019).

### 4.2 Sex differences in sgACC rsFC markers of depression risk and severity

We did not observe sex differences in depression symptom severity, likely because the HCP data reflect a largely non-clinical sample (Miles et al., 2021). Nonetheless, we noted sex differences in sgACC connectivity markers both in association with polygenic risk and depression severity. A recent study of healthy young adults revealed that, in response to neutral faces, women but not men demonstrated PRS-and anhedonia-related attenuation in regional activations spanning the ACC, amygdala, insula, postcentral gyrus, and cerebellum (Mareckova et al., 2020). It remains to be seen whether or how the current findings of sex differences in sgACC connectivities in relation to PRS and depression severity may contribute to the different clinical presentations of depression in men and women (Bangasser and Cuarenta, 2021).

We found that, in men but not in women, greater PRS and higher depression severity was associated with higher sgACC connectivity with the posterior (CBL-IX) and lower sgACC rsFC with anterior cerebellum (CBL-IV), respectively, suggesting that multiple sgACC-cerebellum circuits disposing individuals to depression and manifesting the severity of depression. Meta-analyses of the functional topography as well as human lesion studies support a posterior cognitive/emotional vs. anterior sensorimotor dichotomy of the human cerebellum (Clausi et al., 2019; Stoodley et al., 2016; Stoodley and Schmahmann, 2009). Altered sgACC-posterior cerebellar connectivity may reflect genetically informed neural basis of dysfunctional cognitive/emotional regulation in men. On the other hand, previous studies also emphasized the impact of dysfunctional sensorimotor regulation on mood, with hypo or hyper-activation of the sensorimotor system contributing to the aggravation of depression symptoms (Canbeyli, 2010). Our findings suggest the sgACC-anterior cerebellar rsFC changes in young men may result in the dysregulation of sensorimotor processing and in turn higher severity of depression symptoms.

We also found higher sgACC-occipital cortical coupling in young females showing elevated polygenic risks for depression. Prior research has implicated the occipital cortex in depression (Koch and Schultz, 2014; Liu et al., 2022). An earlier morphometric study showed that larger volumes of the lingual gyrus were associated with preserved cognitive function in MDD patients (Jung et al., 2014). Young adults with a family history of depression vs. control showed higher glutamate/creatine ratios, suggesting metabolic changes, in the occipital cortex (Taylor et al., 2011). Our findings add to this literature and suggest that abnormal fronto-occipital connectivity may reflect genetic vulnerability to depression.

In men but not in women, lower sgACC-insula connectivity was associated with higher depression severity, consistent with the role of frontal-limbic circuit in emotional regulation and depression. A previous study also showed lower sgACC-insular functional connectivity in association with higher depression scores in older healthy adults (mean age 56 years) but did not examine the sex differences (Philippi et al., 2015). Another study demonstrated that greater sgACC-insula connectivity in healthy young men vs. women but did not related sex differences in how sgACC-insula related to any clinical measurements (Wang et al., 2014). With extensive structural and functional connections with the ACC, the insula supports interoception and integrates interoceptive and motivational signals in behavioral control (Namkung et al., 2017). Prior evidence has linked glutamatergic deficits in the patients with MDD to aberrant ACC responses to emotional stimuli and altered ACC-insula connectivity (Horn et al., 2010; Walter et al., 2009). It remains to be seen whether glutamatergic mechanisms might play a sex-specific role in shaping sgACC-insula dysconnectivity in depression. Studies of sgACC connectivities during task challenges, e.g., emotion exposure, regulation, and memory, may help in unraveling the mechanisms underlying sex differences in the clinical manifestations of depression (Bangasser and Cuarenta, 2021; Chaudhary et al., 2023).

Moreover, after controlling for the somatic complaints *T* score in the regression, we identified sgACC-hippocampus rsFC in negative correlation with depression severity in men and sgACC-MTG and sgACC-SMA rsFC each in positive correlation with PRS and depression severity in women. These sgACC rsFC markers may reflect sex-specific markers of depression other than its somatic manifestation. Converging evidence has highlighted the role of hippocampus in emotional regulation and the pathophysiology of depression (Campbell and MacQueen, 2004; Liu et al., 2017) as well as sex-specific hippocampal structural and functional alterations in response to stress (Gobinath et al., 2014; Liu et al., 2019; Naninck et al., 2011). The hippocampal efferents include the prefrontal cortex and ACC (MacQueen and Frodl, 2011). Our findings suggest male-specific disruption of hippocampal-sgACC interaction in association with depression severity. In women alone, we observed higher sgACC-right SMA rsFC in association with greater depression severity. The SMA supports incentivized motor behaviors with and has been linked to psychomotor disturbance in depressive disorders (Exner et al., 2009; Zhang et al., 2016). Our findings suggest that the contribution of SMA dysfunction to depression may be female-specific.

### 4.3 PRS for depression and severity of somatic complaints

We found that, amongst the clinical measures of ASR, higher severity of somatic complaints was significantly related to greater PRS for depression. Indeed, individuals with depression typically report somatic problems, including fatigue, headache, and pain (Kapfhammer, 2006; Sayar et al., 2003; Shim et al., 2020). The effect size of the correlations appeared to be higher between somatic complaints (vs. depression score) and PRS. Thus, the finding suggested somatic complaints rather than depression per se as a more prominent symptom in people at risk for depressive disorder in a neurotypical sample. Along with previous findings of specific SNPs in association with somatic complaints in patients with MDD (Klengel et al., 2011), our findings further linked somatic complaints with overall genetic risks for depression, indicating a shared genetic basis in depression and somatic complaints.

We observed that higher severity of somatic complaints was associated with weaker sgACC rsFC with the left postcentral and supramarginal gyri (PoCG/SMG) across all subjects. This finding broadly aligns with prior evidence linking more severe somatic complaints to diminished gray matter volumes in both the ventromedial prefrontal cortex and somatosensory cortex in non-clinical populations (Wei et al., 2015; Wei et al., 2020). Patients with somatic symptom disorders vs. healthy controls showed decreased sgACC rsFC with the inferior parietal gyrus (Ji et al., 2021), a region encompassing the PoCG and SMG and known for its role in sensory integration and awareness (Teixeira et al., 2014). Rodent studies highlighted the sgACC as a key region of the visceromotor network that modulates autonomic and neuroendocrine responses to stressful stimuli (Drevets et al., 2008). Diminished sgACC connectivity with the sensory and sensory association cortices may contribute to clinical signs and somatic symptoms in mood disorders.

We also found reduced sgACC-dPCu rsFC in correlation with higher severity of somatic complaints in women, although men and women did not differ in the somatic complaint *T* scores. Both dorsal and ventral PCu are involved in processing affective responses to somatic symptoms (Delvecchio et al., 2019; Rossetti et al., 2021). Previous studies identified both structural and functional alterations in the PCu and ACC in individuals with elevated somatic symptoms (Lemche et al., 2013; Su et al., 2014; Szymkowicz et al., 2017). However, few have examined the potential sex-specific neural markers underlying somatic complaints. An earlier study revealed that higher levels of somatic symptoms predicted an increase in proinflammation in female but not male patients with MDD (Dannehl et al., 2014). More research integrating brain markers, immunoreactivity, and clinical symptoms may further unravel sex differences in the pathophysiology of depression.

### 4.4 Limitations and conclusions

The current study is subject to a few limitations. First, the HCP sample represented a largely healthy population without clinical diagnoses. It remains to be seen whether the current findings could be generalized to patients with depressive disorders. Second, except for alcohol use severity, we did not consider other clinical characteristics or comorbidities, such as anxiety, in covariate analyses (Savitz and Drevets, 2009). Future research may examine how comorbidities of depression would influence the neurobiological markers of depression and its genetic risks. Third, the HCP study was cross-sectional, and thus we are not able to investigate the stability of the neural correlates in the manifestation of the genetic risks of depression. Longitudinal data would be most valuable to further distinguish neural phenotypes of genetic risks and clinical severity of depression.

In conclusion, we revealed alterations in the sgACC-DMN and sgACC-frontal connectivity in association with polygenic risks for depression. We also uncovered sex differences in the sgACC functional connectivity implicated in polygenic risks, depression symptoms, and somatic complaints. By integrating neuroimaging and genetics, we can make substantial progress toward understanding the pathophysiology of depression, identifying individuals at risk for early treatments, and establishing reliable benchmarks to predict treatment response.

## CRediT authorship contribution statement

**Yu Chen:** Conceptualization, Methodology, Software, Data Curation, Formal Analysis, Writing – Original Draft, Writing – Review & Editing, Visualization. **Huey-Ting Li:** Software, Data Curation, Formal Analysis, Writing – Original Draft, Writing – Review & Editing. **Xingguang Luo:** Methodology, Software, Data Curation, Formal Analysis, Writing – Original Draft, Writing – Review & Editing. **Guangfei Li:** Formal Analysis, Writing – Original Draft, Writing – Review & Editing. **Jaime S. Ide:** Formal Analysis, Writing – Original Draft, Writing – Review & Editing. **Chiang-Shan R. Li:** Conceptualization, Methodology, Writing – Original Draft, Writing – Review & Editing, Supervision, Funding Acquisition.

## Acknowledgements

This study is supported by NIH grant DA051922. The NIH is otherwise not responsible for the conceptualization of the study, data collection and analysis, or in the decision in publishing the results.

## Competing interests

The authors declare no competing interests in the current study.

## Data availability, Ethics declarations, and Consent to participate

We have obtained permission from the Human Connectome Project (HCP) to use the Open and Restricted Access data for the current study. Data were provided by the WU-Minn Consortium (Principal Investigators: David Van Essen and Kamil Ugurbil; 1U54MH091657) funded by the 16 NIH Institutes and Centers that support the NIH Blueprint for Neuroscience Research; and by the McDonnell Center for Systems Neuroscience at Washington University. The HCP young-adult data is publicly available at https://www.humanconnectome.org/study/hcp-young-adult/.

## Notes

### Competing Interest Statement

The authors have declared no competing interest.

