## Supplementary Materials for "Polygenic risk for depression and resting state functional connectivity of subgenual anterior cingulate cortex in young adults"

**Supplementary Table S1.** The sgACC rsFC correlates of depression *T* score (depr*T*) and polygenic risk score (PRS) for all, men, and women

| Cluster size (*k*) | Voxel *Z*-score | MNI Coordinates (mm) | | | Identified Region |
|---|---|---|---|---|---|
| | | x | y | z | |
| ***All*** | | | | | |
| *deprT* | | | | | |
| 104 | -4.67 | 4 | -46 | -18 | Right cerebellum lobule IV |
| *PRS* | | | | | |
| 281 | -4.60 | -10 | 48 | 42 | Bilateral superior frontal gyri |
| 103 | -4.03 | 4 | -50 | 18 | Bilateral posterior cingulate cortex and ventral precuneus |
| ***Men*** | | | | | |
| *deprT* | | | | | |
| 108 | -4.44 | -36 | 8 | -12 | Left insula |
| 106 | -4.35 | 2 | -60 | -14 | Right cerebellum lobule IV |
| *PRS* | | | | | |
| 117 | 4.18 | -20 | -38 | -48 | Left cerebellum lobule IX |
| 177 | -4.89 | -10 | 48 | 42 | Left superior frontal gyrus |
| ***Women*** | | | | | |
| *deprT* | | | | | |
| Nil | | | | | |
| *PRS* | | | | | |
| 247 | 4.21 | -4 | -72 | 10 | Bilateral lingual gyri and calcarine sulcus |

*Note:* The depr*T* and PRS were modeled together in whole-brain regressions, with age, sex (for all subjects), race, drinking principal component (PC1), and household income as covariates. The results were evaluated at voxel $p < 0.05$ uncorrected in combination with cluster $p < 0.05$ family-wise error (FWE) corrected.

**Supplementary Table S2.** The sgACC rsFC correlates of depression *T* score (depr*T*), polygenic risk score (PRS), and somatic complaints *T* score (*scT*) for all, men, and women

| Cluster size (*k*) | Voxel *Z*-score | MNI Coordinates (mm) | | | Identified Region |
|---|---|---|---|---|---|
| | | x | y | z | |
| ***All*** | | | | | |
| *deprT* | | | | | |

| | | | | | |
|---|---|---|---|---|---|
| 375 | -4.45 | 4 | -46 | -18 | Right cerebellum lobule IV |
| *PRS* | | | | | |
| 277 | -4.55 | -10 | 48 | 42 | Bilateral superior frontal gyri |
| 122 | -4.08 | -4 | -50 | 18 | Bilateral posterior cingulate cortex and ventral precuneus |
| *scT* | | | | | |
| 161 | -4.51 | -54 | -26 | 40 | Left supramarginal gyrus, inferior parietal gyrus, and postcentral gyrus |
| ***Men*** | | | | | |
| *deprT* | | | | | |
| 159 | 4.57 | 28 | 36 | 2 | None |
| 164 | -4.96 | -36 | 8 | -12 | Left insula |
| 136 | -4.28 | -24 | -14 | -10 | Left hippocampus |
| 261 | -4.25 | 2 | -60 | -14 | Bilateral cerebellum lobule IV |
| *PRS* | | | | | |
| 116 | 4.19 | -20 | -38 | -48 | Left cerebellum lobule IX |
| 172 | -4.87 | -10 | 48 | 42 | Left superior frontal gyrus |
| *scT* | | | | | |
| Nil | | | | | |
| ***Women*** | | | | | |
| *deprT* | | | | | |
| 130 | 4.16 | 6 | 10 | 60 | Right supplementary motor area |
| 118 | -4.59 | 18 | -56 | 10 | Right calcarine sulcus |
| *PRS* | | | | | |
| 97 | 4.28 | 46 | -56 | 10 | Right middle temporal gyrus |
| 101 | 4.16 | -4 | -72 | 10 | Bilateral lingual gyri and calcarine sulcus |
| 154 | 4.11 | 6 | -86 | -2 | Bilateral lingual gyri and calcarine sulcus |
| *scT* | | | | | |
| 98 | -4.60 | 2 | -48 | 70 | Bilateral dorsal precuneus |

*Note:* The depr*T*, PRS, and sc*T* were modeled together in whole-brain regressions, with age, sex (for all subjects), race, drinking principal component (PC1), and household income as covariates. The results were evaluated at voxel $p < 0.05$ uncorrected in combination with cluster $p < 0.05$ family-wise error (FWE) corrected.

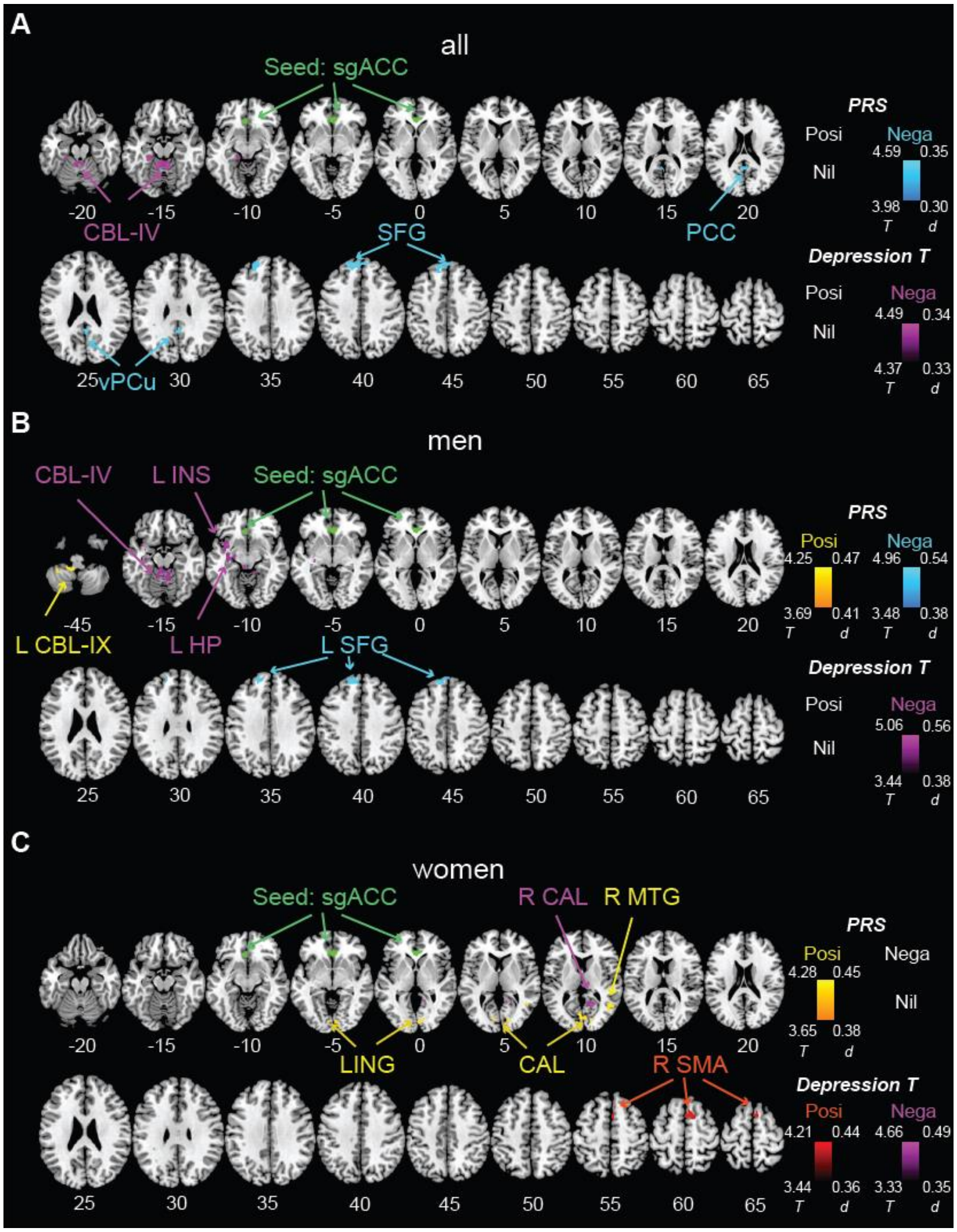

**Supplementary Figure S1.** Seed-based whole-brain rsFC of sgACC (green) in correlation with polygenic risk score (PRS) and depression *T* score for **(A)** all subjects, **(B)** men, and **(C)** women, when PRS, depression *T* score, and somatic complaints *T* score were modeled together, with age, sex (for all subjects), race, drinking principal component (PC1), and household income as covariates. The results were evaluated at voxel $p < 0.001$, uncorrected in combination with cluster $p < 0.05$ family-wise error (FWE) corrected. Warm color: in positive (Posi) correlation with PRS; Cool color: in negative (Nega) correlation with PRS; Red: in positive (Posi) correlation with depression *T* score; Violet: in negative (Nega) correlation with depression *T* score; Color bar shows voxel *T* and Cohen's *d* values. L: left; R: right; CBL: cerebellum; sgACC: subgenual anterior cingulate cortex; PCC: posterior cingulate cortex; vPCu: ventral precuneus; SFG: superior frontal gyrus; INS: insula; LING: lingual gyrus; CAL: calcarine sulcus; MTG: middle temporal gyrus; SMA: supplementary motor area.
